# Visual adaptation and microhabitat choice in Lake Victoria cichlid fish

**DOI:** 10.1101/443655

**Authors:** Daniel Mameri, Corina van Kammen, Ton G.G. Groothuis, Ole Seehausen, Martine E. Maan

**Affiliations:** CEF – Forest Research Centre, Instituto Superior de Agronomia, University of Lisbon, Lisboa, Portugal; Groningen Institute for Evolutionary Life Sciences (GELIFES), University of Groningen, Groningen, The Netherlands; Van Hall Larenstein University of Applied Sciences, Leeuwarden, The Netherlands; Department of Fish Ecology & Evolution, Eawag, Center for Ecology, Evolution and Biogeochemistry, Kastanienbaum, Switzerland; Institute of Ecology & Evolution, University of Bern, Bern, Switzerland

**Keywords:** sensory drive, habitat choice, haplochromine, ecological speciation, colour vision

## Abstract

When different genotypes choose different habitats to better match their phenotypes, adaptive differentiation within a population may be promoted. Mating within those habitats may subsequently contribute to reproductive isolation. In cichlid fish, visual adaptation to alternative visual environments is hypothesised to contribute to speciation. Here, we investigated whether variation in visual sensitivity causes different visual habitat preferences, using two closely related cichlid species that occur at different but overlapping water depths in Lake Victoria and that differ in visual perception (*Pundamilia* sp.). In addition to species differences, we explored potential effects of visual plasticity, by rearing fish in two different light conditions: broad-spectrum (mimicking shallow water) or red-shifted (mimicking deeper waters). Contrary to expectations, fish did not prefer the light environment that mimicked their typical natural habitat. Instead, we found an overall preference for the broad-spectrum environment. We also found a transient influence of the rearing condition, indicating that assessment of microhabitat preference requires repeated testing to control for familiarity effects. Together, our results show that cichlid fish exert visual habitat preference, but do not support straightforward visual habitat matching.

## Introduction

In heterogeneous environments, individuals may disperse to (micro)habitats that best match their phenotype - i.e. ‘matching habitat choice’ [1]. This behaviour may dissipate natural selection for local adaptation but, if it causes habitat segregation, it may contribute to genetic differentiation between (micro)habitats and ultimately speciation [2-3]. Here, we investigate this process in the context of sensory drive, testing whether divergent visual phenotypes preferentially seek out alternative visual environments.

Visual systems adapt rapidly, responding to environmental challenges associated with foraging, predator avoidance and (sexual) communication [4-6]. This is particularly well-documented in visually heterogeneous aquatic environments [7-9]. In Lake Victoria (East Africa), divergent visual adaptation is associated with speciation in the genus *Pundamilia* [6,10,11]. Replicate pairs of sympatric species consist of one species with blue male coloration (*P. pundamilia* and P. sp. *‘pundamilia-like’*) and one with red/yellow male coloration (P. *nyererei* and *P. sp. ‘nyererei-like’* [11]). The species inhabit different (but overlapping) depth ranges and thereby experience different light environments: ‘blue species’ tend to inhabit shallow waters, receiving broad-spectrum light, while ‘red species’ tend to inhabit deeper waters with red-shifted light conditions [6]. They differ in opsin gene sequence (light-sensitive proteins in the eye; [6]) and opsin gene expression [12], and in visual response to blue and red light [9], corresponding to the difference in visual habitat. Recent work suggests that at least some of these differences are adaptive: when raising the fish in artificial light conditions that mimic shallow and deep habitats, both species survive best in their own natural light condition [13].

In this study, we test whether differences in visual sensitivity between P. *sp. ‘pundamilia-like’* and *P. sp. ‘nyererei-like’* cause different visual habitat preferences. We expect that when given a choice, individuals will disperse from a suboptimal visual environment to one that better matches their visual system phenotype [1]. In addition to genetic differences, developmental plasticity may contribute to variation in visual sensitivity [14]. To explore this, and to assess the causal relationship between visual sensitivity and habitat preference, we manipulate visual development by raising the fish under different light conditions.

We predict that individuals of either species prefer the light regime that is closest to the one their populations are adapted to. First-generation interspecific laboratory-bred hybrids are expected to have no preference, because their visual system presumably has intermediate characteristics, and they survive equally well in both environments [13]. We also predict that fish prefer the light environment they are reared in, due to environment-induced plastic changes in visual sensitivity, particularly in hybrids that may lack genetically determined visual specialization.

## Material and Methods

### Fish

We used F1 and F2 sub-adults, bred in captivity from wild-caught P. *sp. ‘pundamilia-like’* and *P. sp. ‘nyererei-like’* from Python Islands [6]. Fish were maintained in family groups, divided equally over two light treatments (details in Supplementary Materials) that mimicked the natural light environments experienced by *P. sp. ‘pundamilia-like’* (shallow water, 0-2 m) and *P. sp. ‘nyererei-like’* (deeper water, 0-5 m) at Python Islands [6]. Fish were tested in groups of fixed composition (rather than individually, to minimise stress), with 4 siblings from the same light treatment. We used 5 sibling groups for each species and for the hybrids, from each condition, generating a total of 30 groups. Fish were not individually recognized and only group-level data was recorded. Until testing, fish were naïve to the light condition they were not reared in.

### Experimental setup and procedures

The experimental tank (112×46×41 cm) was divided into two equally-sized compartments by an opaque PVC sheet, with a semi-circular hole of 10 cm diameter at the bottom to allow movement between sides (Figures S4-S5). One side of the tank had the shallow light condition (broad-spectrum) and the other one the deep light condition (red-shifted spectrum), which could be reversed.

For 1 hour, we recorded the number of fish on each side. As a measure of activity, we also counted the number of times individuals crossed between sides. Trials were considered successful if at least four crossings were recorded. Groups were excluded if unsuccessful twice.

All groups were tested twice, with ~2 weeks between trials. Light environments were switched between tank sides after the first trial. After analysing the first two trials, we submitted *P. sp. ‘pundamilia-like’* and *P. sp. ‘nyererei-like’* to a third trial to increase statistical power for testing species differences.

### Data analyses

We calculated the proportion of time spent on the side of the tank with shallow light conditions (summed for the individuals in a group) as a measure of preference, ranging from 0 to 1. We used linear mixed effects models with arcsine-transformed preference scores (R 3.5.1 [15]; packages ‘lme4’ and ‘lmerTest’). Fixed factors included species (*P. sp. ‘pundamilia-like’, P. sp. ‘nyererei-like’* and hybrids), rearing environment, activity and age. Random effects included trial number, nested in fish group, nested in family (some groups came from the same family – see Table S1). We selected the minimum adequate models (lowest AIC) that significantly differed from the null model (Satterthwaite’s ANOVA).

## Results

75 of the 80 trials were successful. Overall, fish preferred the blue-shifted (shallow) light environment (Figure 1; mean±SE: 61±21%; z=4.086, p<0.001). They also showed a significant preference for the environment they were reared in (F_1,42.638_=6.9474, p=0.012). However, the difference between rearing groups was only seen in the first trial: shallow-reared fish expressed a consistent preference for the shallow environment, while deep-reared fish initially had no preference but developed a preference for shallow in subsequent trials (trial effect in deep-reared fish: F_1,23.618_=7.3824, p=0.01213; interaction between rearing environment and trial: F_1,_ 47.6=3.0567, p=0.0869).

**Figure 1.**
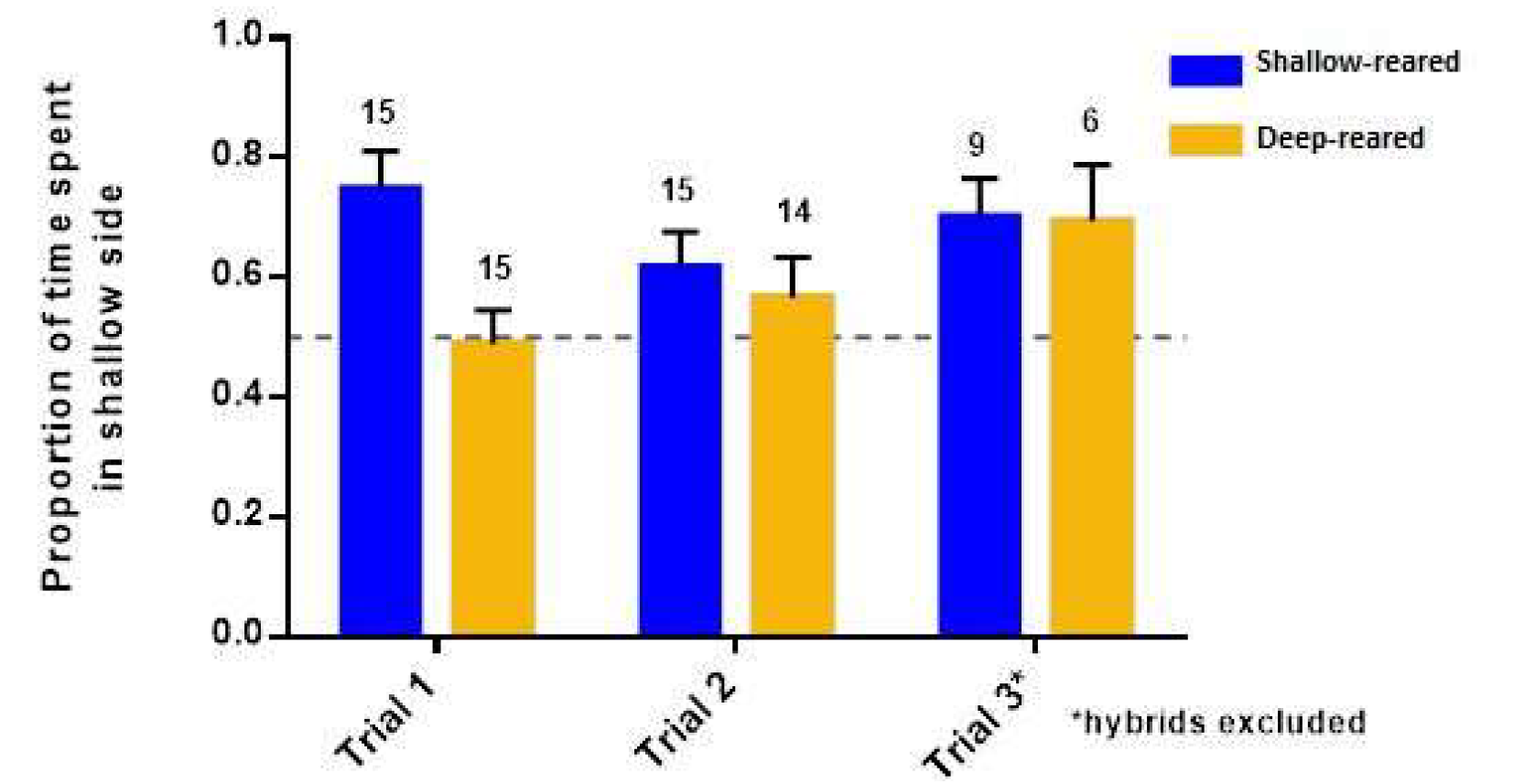
Fish spent more time in the shallow light condition, but deep-reared fish expressed no preference in the first trial. Bars are means with standard errors; Blue bars: shallow-reared fish; yellow bars: deep-reared fish. Numbers above bars indicate the number of test groups.

There was no difference in preference between the three genetic groups (F_2,8.468_=0.6044, p=0.57; Figure 2). Repeating the analyses without hybrids also did not reveal differences between *P. sp. ‘pundamilia-like’* and *P. sp. ‘nyererei-like’* groups (F_1, 6.6704_=3.1233, p=0.12).

**Figure 2.**
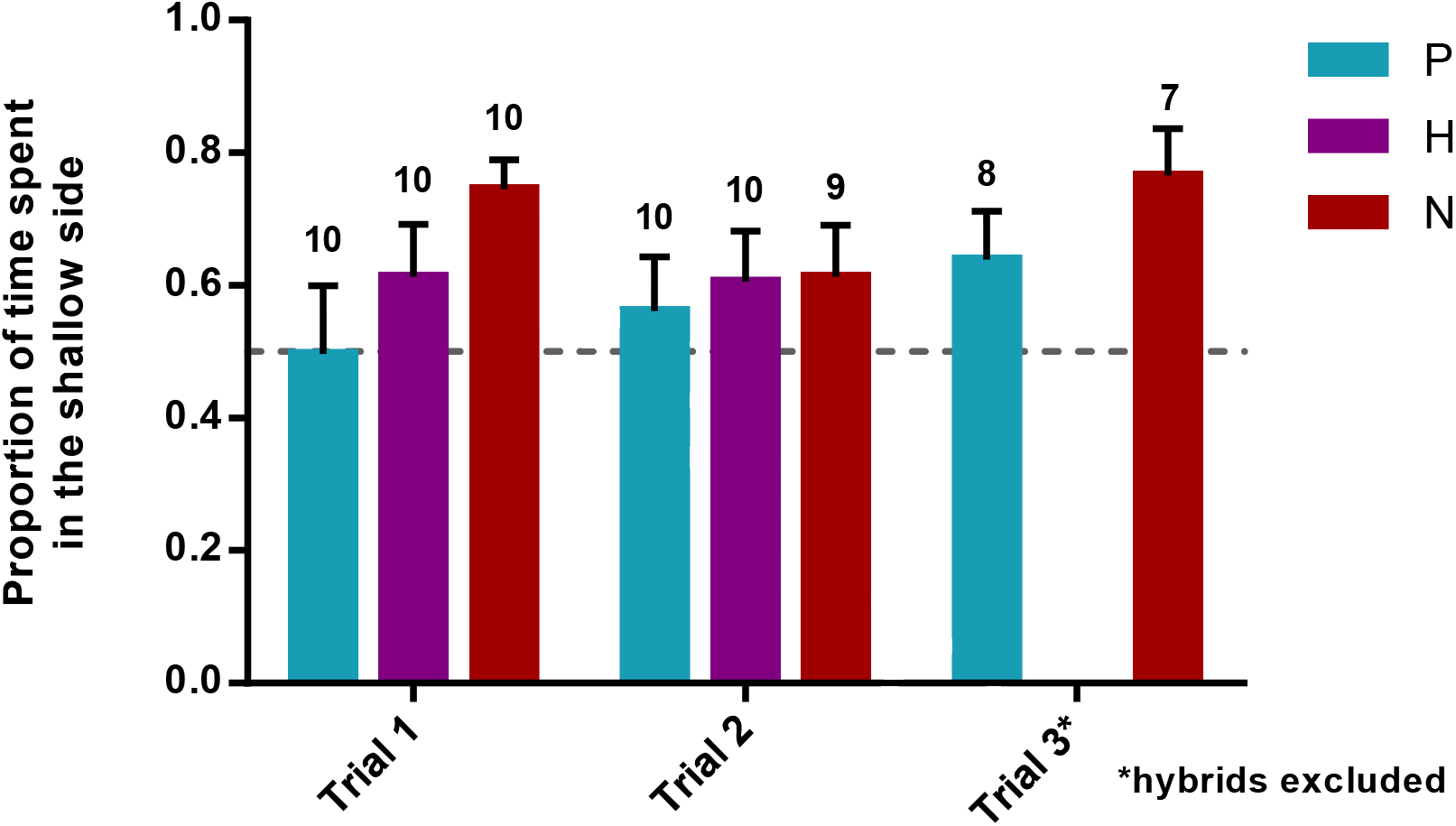
Visual habitat preference in *P. sp. ‘pundamilia-like’* (‘P’, blue), hybrids (‘H’, purple) and *P. sp. ‘nyererei-like’* (‘N,’ red), in trials 1, 2 and 3. Bars are means with standard errors; numbers above bars indicate the number of test groups.

The minimal adequate model explaining preference also included fish activity (mean±SE=34.6±3.37 crossings per trial): more active groups expressed weaker preferences for the shallow light condition (F_1, 69.742_=8.383, p=0.005). To explore this further, we calculated preference strength (deviation from 0.5, irrespective of the chosen light condition) and found that more active groups expressed weaker preferences overall (F_1, 71.837_=14.945, p=0.0002; Figure S8). Fish activity did not significantly differ between species (F_2,7.2456_=3.1968, p=0.1011), rearing conditions (F_1,67.317_= 0.3673,p=0.5465) nor trials (F_1, 56.345_ =0.2172, p=0.643).

## Discussion

Matching habitat choice can evolve in response to selection for improving performance in heterogeneous environments [1]. Here, we investigated this phenomenon in two closely related cichlid species with divergent visual system characteristics, testing the hypothesis that individuals should preferentially reside in the light environment that mimics their natural habitat.

Contrary to predictions, the shallow water-dwelling *P. sp. ‘pundamilia-like’* and the deeper-dwelling *P. sp. ‘nyererei-like’* did not differ in visual habitat preference. Instead, we found an overall preference for the broad-spectrum light condition, mimicking shallow waters. This is surprising, given that opsin genotype is subject to divergent selection between these species, as evidenced by signatures of divergent selection (on the long-wavelength-sensitive opsin gene, LWS [6,11]) and differences in survival between light environments in captivity [13]. In other fish, preferences for light conditions that maximize performance have been demonstrated [16,17]. Possibly, our fish did not have enough opportunity to evaluate their performance in the two environments: while we added food cues to stimulate exploration, we did not provide an actual reward. Matching habitat choice may be most pronounced when it generates substantial reward [18].

We found a transient effect of the rearing light environment: in the first trial, deep-reared fish did not express a preference for the shallow light condition. Developmental effects on visual habitat preference have been observed in some fish species but not others (e.g. Australasian snapper prefer light intensities that match their rearing environment [16], but Coho salmon prefer darker backgrounds even when raised in bright illumination [19]). In the present study, it seems that familiarity with the rearing environment may have suppressed exploration of the unfamiliar one [17]. To explore this further, we also assessed preference in 15-minute blocks within the first trial. We did not observe that deep-reared fish gradually spent more time in the unfamiliar environment (shallow) in the course of this first trial, suggesting that habituation requires longer or more frequent exposure (Figure S7). Either way, this finding entails a warning for future studies: testing individuals only once may poorly estimate behavioural preferences.

To conclude, we find that *Pundamilia* cichlid fish exert significant preference for visual habitat, preferring broad-spectrum over red-shifted light conditions. Species differences in visual traits do not translate into differences in preference. Light conditions during development do influence preference, but only in the short term. We conclude that our results do not support a simple role of vision-mediated matching habitat choice in *Pundamilia* cichlids.

## Authors’ contributions

MEM, OS and TGG designed the study; DM and MEM developed the experimental setup; DM and CvK collected the data. DM analysed the data; DM and MEM wrote the manuscript, with contributions from OS and TGG. All authors gave final approval for publication.

## Acknowledgements

We acknowledge the Tanzanian Commission for Science and Technology for research permission and the Tanzanian Fisheries Research Institute for hospitality and facilities. We thank M. Haluna, M. Kayeba, E. Ripmeester, O. Selz for help in the field, S. Veenstra, B. Verbeek, A. Taverna and E. Schaeffer for taking care of the fish in the laboratory, and S. Wright and G. Nunes for help with fish care and experimental design.

## References

1. Edelaar P, Siepielski AM, Clobert J. 2008 Matching habitat choice causes directed gene flow: aneglected dimension in evolution and ecology. Evolution 62, 2462–2472. (doi:10.1111/j.1558-5646.2008.00459.x)

2. Edelaar P, Bolnick DI. 2012 Non-random gene flow: an underappreciated force in evolution and ecology. Trends in Ecology & Evolution 27, 659–665. (doi:10.1016/j.tree.2012.07.009)

3. Jiang Y, Peichel CL, Torrance L, Rizvi Z, Thompson S, Palivela VV, Pham H, Ling F, Bolnick DI. 2017 Sensory trait variation contributes to biased dispersal of threespine stickleback in flowing water. Journal of Evolutionary Biology 30, 681–695. (doi:10.1111/jeb.13035)

4. Endler JA. 1992 Signals, Signal Conditions, and the Direction of Evolution. The American Naturalist 139, S125–S153. (doi:10.1086/285308)

5. Stevens M. 2013 Sensory Ecology, Behaviour and Evolution. Oxford University Press, UnitedKingdom, 264 pp.

6. Seehausen O, Terai Y, Magalhães IS, Carleton KL, Mrosso HDJ, Miyagi R, van der Sluijs I, Schneider MV, Maan ME, Tachida H, Imai H, Okada N. 2008 Speciation through sensory drive in cichlid fish. Nature 455, 620–626. (doi:10.1038/nature07285)

7. Fuller RC, Noa LA. 2010 Female mating preferences, lighting environment, and a test of the sensory bias hypothesis in the bluefin killifish. Animal Behaviour 80, 23–35. (doi:10.1016/j.anbehav.2010.03.017)

8. Boughman JW. 2002 How sensory drive can promote speciation. Trends in Ecology & Evolution 17, 571–577. (doi:10.1016/s0169-5347(02)02595-8)

9. Maan ME, Hofker KD, van Alphen JJM, Seehausen O. 2006 Sensory Drive in Cichlid Speciation. The American Naturalist 167, 947–954. (doi:10.1086/503532)

10. Meier JI, Marques DA, Mwaiko S, Wagner CE, Excoffier L, Seehausen O. 2017 Ancient hybridization fuels rapid cichlid fish adaptive radiations. Nature Communications 8, 14363. (doi:10.1038/ncomms14363).

11. Meier JI, Marques DA, Wagner CE, Excoffier L, Seehausen O. 2018 Genomics of Parallel Ecological Speciation in Lake Victoria Cichlids. Molecular Biology and Evolution msy051. (doi:10.1093/molbev/msy051)

12. Carleton KL, Parry JWL, Bowmaker JK, Hunt DM, Seehausen O. 2005 Colour vision and speciation in Lake Victoria cichlids of the genus Pundamilia. Molecular Ecology 14, 4341–4353. (doi:10.1111/j.1365-294x.2005.02735.x)

13. Maan ME, Seehausen O, Groothuis TGG. 2017 Differential Survival between Visual Environments Supports a Role of Divergent Sensory Drive in Cichlid Fish Speciation. The American Naturalist 189, 78–85. (doi:10.1086/689605)

14. Wright DS, Demandt N, Alkema JT, Seehausen O, Groothuis TGG, Maan ME. 2016 Developmental effects of visual environment on species-assortative mating preferences in Lake Victoria cichlid fish. Journal of Evolutionary Biology 30, 289–299. (doi:10.1111/jeb.13001)

15. R Core Team. 2018. R: A language and environment for statistical computing. R Foundation for Statistical Computing, Vienna, Austria. URL: https://www.R-project.org/

16. Robinson E, Jerrett A, Black S, Davison W. 2017 Snapper rest where they see best: visually mediated choice behaviour of Australasian snapper (*Chrysophrys auratus*). Marine and Freshwater Behaviour and Physiology 50, 81–88. (doi:10.1080/10236244.2017.1304154)

17. Davis J. 2004 The effect of natal experience on habitat preferences. Trends in Ecology & Evolution 19, 411–416. (doi:10.1016/j.tree.2004.04.006)

18. Benkman CW. 2016 Matching habitat choice in nomadic crossbills appears most pronounced when food is most limiting. Evolution 71, 778–785. (doi:10.1111/evo.13146)

19. Gaffney LP, Franks B, Weary D, von Keyserlingk MAG. 2016. Coho salmon (*Oncorhynchus kisutch*) prefer and are less aggressive in darker environments. PLoS One 11, e0151325. (doi:10.1371/journal.pone.0151325).

